# An offset ON-OFF receptive field is created by gap junctions between distinct types of retinal ganglion cells

**DOI:** 10.1101/2020.07.15.205336

**Authors:** Sam Cooler, Gregory W. Schwartz

**Affiliations:** Northwestern University Interdepartmental Neuroscience Graduate Program; Departments of Ophthalmology and Physiology, Feinberg School of Medicine, Northwestern University; Department of Neurobiology, Weinberg School of Arts and Sciences, Northwestern University

## Abstract

Receptive fields (RFs) are a foundational concept in sensory neuroscience. The RF of a sensory neuron is shaped by the properties of its synaptic inputs from connected neurons. In the early visual system, retinotopic maps define a strict relationship between the location of a cell’s dendrites and its RF location in visual space^1–3^. Retinal ganglion cells (RGCs), the output cells of the retina, form dendritic mosaics that tile retinal space and have corresponding RF mosaics that tile visual space^1,2^. The precise location of dendrites in some RGCs has been shown to predict their RF shape^4^. Previously described ON-OFF RGCs have aligned dendrites in ON and OFF synaptic layers, so the cells respond to increments and decrements of light at the same locations in visual space^5–8^. Here we report a systematic offset between the ON and OFF RFs of an RGC type. Surprisingly, this property does not come from offset ON and OFF dendrites but instead arises from electrical synapses with RGCs of a different type. This circuit represents a new channel for direct communication between ON and OFF RGCs. Using a multi-cell model, we find that offset ON-OFF RFs could improve the precision with which edge location is represented in an RGC population.

## Introduction

At the first synapse in the visual system, the output of the photoreceptors, signals diverge into ON pathways, which respond to increments of light, and OFF pathways, which respond to decrements of light. ON and OFF pathways reconverge in multiple locations, including at the retinal output in ON-OFF retinal ganglion cells (RGCs). In the mouse, where they are best characterized, RGCs comprise greater than 40 functionally, morphologically, and transcriptomically distinct types^9–14^. All previously identified ON-OFF RGCs share two common features: (1) aligned ON and OFF RFs, and (2) excitatory synaptic inputs from both ON and OFF bipolar cells. Inputs from ON and OFF bipolar cells are formed either at two distinct dendritic strata in the inner plexiform layer (IPL)^8^ or at a single stratum in the middle of the IPL where ON and OFF bipolar cell terminals overlap^15^.

We report a systematic spatial offset between the ON and OFF RF of an RGC type. Instead of a misalignment in its dendrites, we reveal that this RF offset arises from a novel circuit composed of gap junctions with several RGCs of a different type. While RFs with offset ON and OFF subfields result in a modest amount of direction selectivity and orientation selectivity for certain stimuli at the level of single RGCs, modeling demonstrates a large enhancement in the encoding of edge position within a population of RGCs. Our multi-cell model reveals that offset ON and OFF RF sub-fields could help a population of RGCs encode edge position with precision down to 0.6 degrees of visual angle, less than 12% of the RF diameter of a single RGC.

## Results

F-mini RGCs were recently identified as two different cell types: F-mini-ON and F-mini-OFF, based on their expression patterns of several transcription factors, their unique morphologies, and their light responses^16^. F-mini RGCs are the second and third most numerous RGC types in the mouse retina, together comprising 13% of RGCs^13^. We recorded light responses from functionally-identified F-mini RGCs in dark-adapted mouse retina (see Methods) and later confirmed their identity by morphological analysis (Fig. 1a-c) and immunohistochemistry (IHC) (Extended Data Fig. 1). Unlike in the initial report, we found that both F-mini RGC types responded to both light increments (ON) and decrements (OFF) for small spots and moving bars (Fig. 1d-g, Extended Data Fig. 2a,b). We focused primarily on F-mini-ON RGCs in search of a circuit mechanism for the ON-OFF responses.

**Figure 1.**
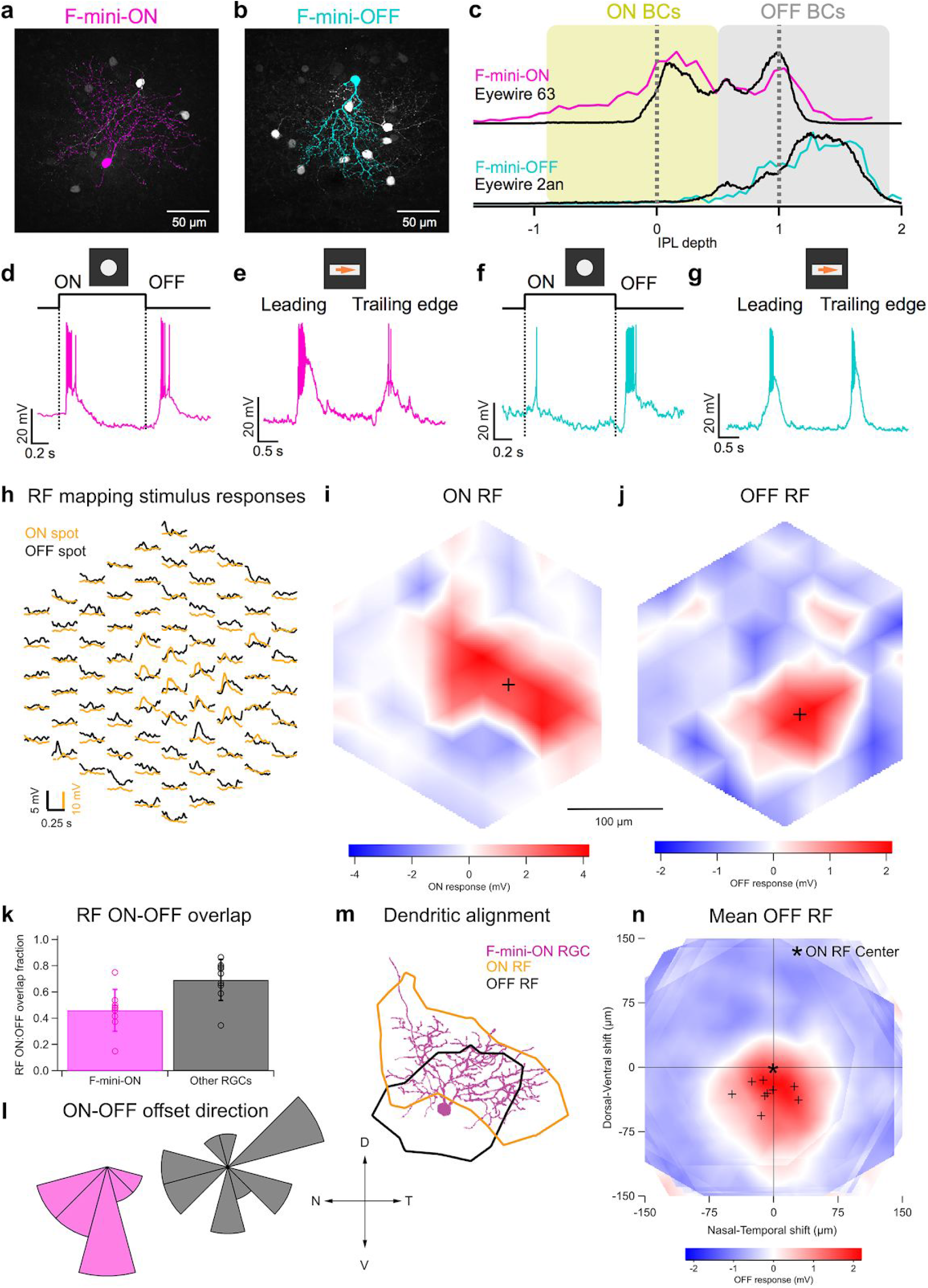
F-mini-ON and F-mini-OFF RGCs have both ON and OFF light responses. **a,b,** Images of F-mini-ON and F-mini-OFF RGCs from fixed tissue. Magenta and cyan color scheme for RGC types is consistent throughout. All cell images throughout are shown with the dorsal direction on the retina toward the top of the page. Targeted RGCs were traced and colored, overlaid on gray. **c,** Stratification profiles of an F-mini-ON and an F-mini-OFF RGC from our cell fills (colored lines) and from the data in the EyeWire museum (black). Dashed lines indicate ON and OFF choline acetyltransferase (ChAT) bands. Shading shows the stratification regions of ON (yellow) and OFF (grey) bipolar cells (BCs) in the inner plexiform layer (IPL). **d,** F-mini-ON RGC in current clamp responding to the onset and offset of a positive contrast spot from darkness to 200 isomerizations (R*)/rod/s. **e,** F-mini-ON RGC responding to the leading and trailing edge of a positive contrast moving bar (140 μm x 500 μm, 1000 μm/s, 200 R*/rod/s). **f, g,** Same as **d, e** for an F-mini-OFF RGC. **h,** Average responses of a F-mini-ON RGC to 30 μm spots of positive (orange) or negative (black) contrast at the indicated positions. **i, j,** Interpolated ON and OFF spatial RFs from the data in **h**. Black crosses mark the center-of-mass (COM). **k,** Population data showing the fractional overlap of the OFF RF relative to the ON RF for F-mini-ON RGCs (magenta) and other ON-OFF RGCs (grey) (see Methods). Points are individual cells and error bars are SD across cells. **l,** Polar histograms showing the offset angle between the ON RF COM and the OFF RF COM for F-mini-ON RGCs (magenta) and other ON-OFF RGCs (grey). OFF ventral of ON is shown as a downward (ventral) angle. **m,** Image of the cell from **h-j** overlaid with its ON (orange) and OFF (black) RF contours. **n,** Average OFF RF for F-mini-ON RGCs aligned to the center-of-mass of each ON RF at the origin. Points are the center-of-mass for individual cells (*n* = 9). Scale bar is the same in **h,i,j,m,n**.

To explore the robustness of the ON-OFF responses in F-mini-ON RGCs, we adapted the retina to different mean luminances across the range from scotopic to photopic. We found robust ON and OFF responses across this range (Extended Data Fig. 2c). We also measured the contrast response functions of F-mini-ON RGCs on a photopic background and found similar contrast sensitivity for the ON and OFF responses (Extended Data Fig. 2d).

We mapped ON and OFF RFs in F-mini RGCs using a stimulus consisting of small spots of positive and negative contrast^7^ (Fig. 1h-j, Extended Data Fig. 3). Surprisingly, we found a consistent dorsal-ventral displacement between the ON and OFF sub-regions of the RF (Fig. 1i-n). In F-mini-ON RGCs we measured a spatial offset of 38 ± 14 µm (36 ± 13% of ON RF diameter; *n* = 9). The vertical component of the offset was always ventralward (Fig. 1n; OFF ventral to ON, −30 ± 13 µm, p < 10^−4^), with a horizontal component not significantly distributed to one side (−7.5 ± 24 µm, p = 0.37). Control ON-OFF RGCs had smaller offset (24 ± 14% of ON RF diameter) and lacked a systematic vertical or horizontal displacement (*n* = 14) (V: −4.0 ± 18 µm, p = 0.42, H: −3.7 ± 23 µm, p = 0.56). The distribution of offset directions was uniform for control ON-OFF RGCs (p = 0.93, Rayleigh test) and highly non-uniform for F-mini-ON RGCs (p < .001, Fig. 1l). We quantified the fractional overlap between the ON and OFF RFs and found it to be lower in F-mini-ON RGCs than in other ON-OFF RGCs (Fig. 1k, p < 0.01). A smaller sample of F-mini-OFF RFs also showed diffuse ON and ventrally displaced OFF sub-fields (Extended Data Fig. 3). Despite this offset between ON and OFF RFs, the population of F-mini-ON RGCs reconstructed in the Eyewire Museum^13^ showed well aligned dendritic strata in the inner and outer half of the IPL, similar to other bistratified RGCs (Extended Data Fig. 4). Thus, F-mini-ON RGCs have RFs with spatially offset ON and OFF sub-regions, with the OFF sub-field consistently ventral of the ON sub-field, and this result cannot be explained by displacement of their dendrites.

Many RGCs make gap junctions with other RGCs of the same type or with amacrine cells^17^, and these electrical synapses can affect RF properties^18,19^. Since dendritic morphology alone could not explain the ON-OFF RF offset in F-mini-ON RGCs, we sought to test whether gap junctions contributed to this unusual RF property. We filled individual F-mini-ON RGCs with the gap-junction permeable tracer Neurobiotin, delivered during whole-cell patch clamp recordings. In addition to the patched cell, Neurobiotin labelled several surrounding cells in the ganglion cell layer with a distinct morphology (Fig. 2a). We confirmed that the coupled cells were RGCs by the presence of an axon extending toward the optic nerve, and we confirmed their identity as F-mini RGCs by their morphology and by the presence or absence of the transcription factors FoxP2 and FoxP1 (Extended Data Fig. 1). Gap junctions between F-mini RGCs were not only permeable to Neurobiotin, but also to the larger fluorescent molecule Alexa Fluor 488 (Fig. 2b). This dye allowed us to identify coupled cells in live tissue by two-photon (2P) excitation^20^, and record their light responses sequentially or simultaneously (Fig. 2c). F-mini-ON RGCs were coupled to 3.85 ± 1.3 RGCs (*n* = 13); F-mini-OFF RGCs to 4.25 ± 2.5 RGCs (*n* = 4, Fig. 2d). In every case in which we attempted to classify a cell directly coupled to an F-mini RGC it was an F-mini RGC of the *other* type. For F-mini-ON RGC injections, all coupled cells tested by IHC were FoxP1+/FoxP2+, the molecular identity of F-mini-OFF RGCs (*n* = 19). All coupled cells tested in live retina had the morphological signature (ventrally directed, compact, OFF-stratifying dendrites) and/or the physiological signature (transient OFF or ON-OFF response with strong surround suppression) of F-mini-OFF RGCs^16^ (*n* = 54). Similar experiments in F-mini-OFF RGCs revealed F-mini-ON RGCs verified by IHC as FoxP1-/FoxP2+ (*n* = 14) or by physiology and morphology (*n* = 24). Regions of dendritic contact between F-mini-ON and F-mini-OFF RGCs did not show immunoreactivity for antibodies against three connexin proteins known to exist in the inner retina, Cx36, Cx45, and Cx30.2 (Extended Data Fig. 5), so the identity of the connexin at these gap junctions remains an open question.

**Figure 2.**
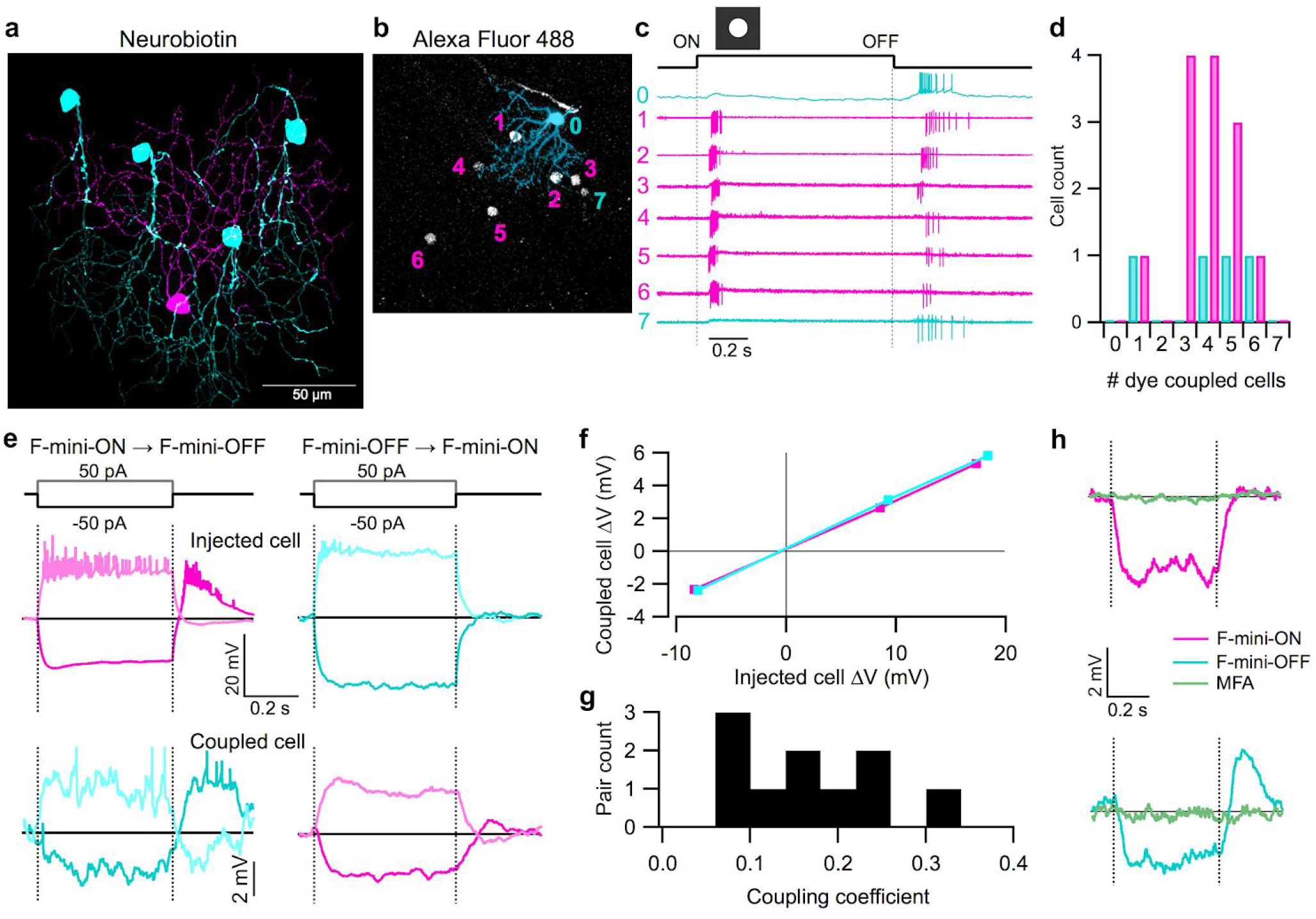
F-mini-ON and F-mini-OFF RGCs are coupled to each other by gap junctions. **a,** Tracing from an experiment in which a single F-mini-ON RGC (magenta) was filled with Neurobiotin. Four coupled F-mini-OFF RGCs are in cyan, and an additional unidentified RGC (possibly F-mini-ON) is in yellow. **b,** The F-mini-OFF RGC labeled ‘0’ was filled with Alexa Fluor 488 (cyan), revealing 7 coupled somas (white). **c,** Cell-attached recordings from each of the labeled somas shown in **b**. Cell 7 is a dimmer, likely a second-order connected F-mini-OFF RGC. **e,** Average voltage traces from a pair in which one F-mini RGC was injected with current (top row) and the coupled F-mini RGC of the opposite type (bottom row) showed a response. Current injections were +50 pA (lighter traces) and −50 pA (darker traces). **f,** Voltage change relationship across the electrical synapse in both directions for the pair in **e**. **g,** Histogram of the coupling coefficient (slope of line in **f**) for all recorded pairs. **h,** Voltage in F-mini RGCs (for −50 pA injection in coupled cell) in control conditions and in the presence of MFA (green).

Having found a dense heterotypically coupled network of RGCs, we sought to test for functional connectivity. We performed paired whole-cell current clamp recordings aided by 2P visualization of Alexa Fluor 488. Both depolarizing and hyperpolarizing current injections were transmitted between the coupled cells, and the resulting voltage transfer was symmetric, the hallmarks of a non-rectifying electrical synapse^21^ (Fig. 2d-f, p > 0.19 for both hyperpolarizing vs. depolarizing current, and injections into F-mini-ON vs. F-mini-OFF RGCs). Coupling coefficient expresses the fraction of the voltage change in one cell that is transmitted to the coupled cell. We measured a coupling coefficient of 0.14 ± 0.08 (range 0.05 – 0.31, *n* = 12) between F-mini-ON and F-mini-OFF RGCs (Fig. 2g), comparable to the strongest coupling coefficients reported in the inner retina, between amacrine cells^22^ (0.25) or between RGCs of the same type^7^ (0.14). Pharmacological block of gap junctions with meclofenamic acid (MFA)^23^ decreased the coupling coefficient to 0.03 ± 0.03 (*n* = 4, Fig. 2h). Additional evidence for functional connectivity between F-mini-ON and F-mini-OFF RGCs came from the fact that their positive noise cross-correlations were proportional to coupling coefficient and decreased by MFA (R^2^ = 0.79, p = 0.007, Extended Data Fig. 6f,i). Thus, F-mini-ON and F-mini-OFF RGCs are not only dye-coupled, but they are also capable of passing substantial amounts of current through their gap junctions, which could potentially mix ON and OFF pathways directly at the level of the RGCs.

To determine how gap junctions with F-mini-OFF RGCs affect the light responses in F-mini-ON RGCs, we sought to measure light responses from the same F-mini-ON RGCs with and without coupling. We used two different manipulations to uncouple F-mini-ON RGCs from their electrical network: pharmacological gap junction block with MFA and physical ablation of coupled RGCs.

We recorded from F-mini-ON RGCs in current-clamp mode and stimulated the retina with positive and negative contrast spots (Fig. 3). MFA eliminated OFF spikes and reduced subthreshold OFF depolarization to 10 ± 10% of control, while only reducing ON spiking to 71 ± 11% and ON depolarization to 97 ± 17% of control (Fig. 3c,d; *n* = 4 spiking, 6 depolarization). The modest reduction in ON spike count despite an unchanged light-evoked depolarization can be attributed to a hyperpolarizing baseline shift (from −61 ± 2.4 mV in control to −65 ± 4.4 mV in MFA, p = 0.05), likely caused by nonspecific effects of MFA^24,25^. Non-F-mini RGCs showed a moderate reduction of spiking in MFA, consistent with reduced contrast sensitivity^24^, but the ON and OFF pathways were affected similarly (Extended Data Fig. 7).

**Figure 3.**
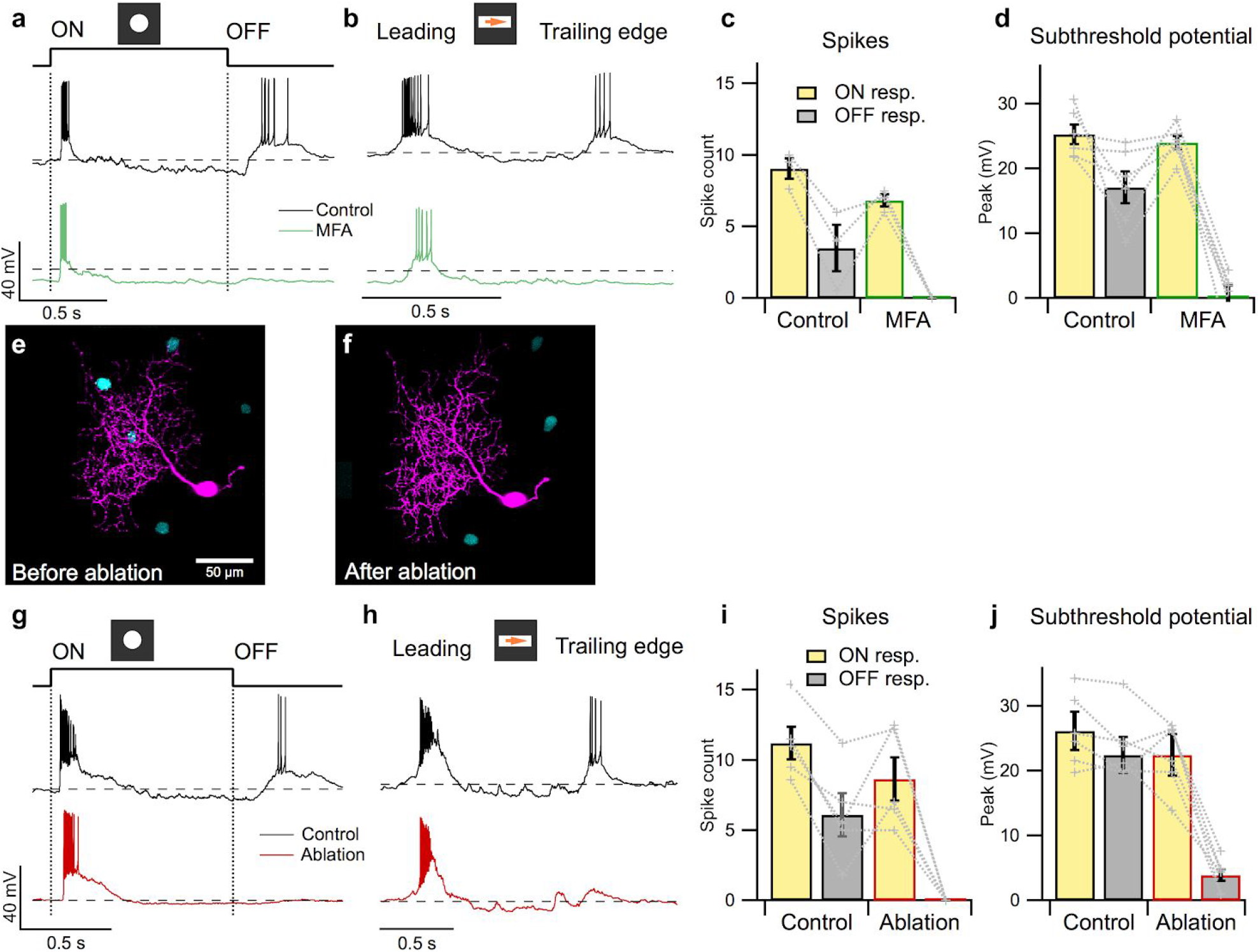
F-mini-ON RGCs receive ON input from chemical synapses and OFF input from electrical synapses. **a,** Response of an F-mini-ON RGCs to an ON 120 µm spot before (black) and after MFA (green) application. Dashed line is −60 mV. **b,** Responses of the cell in **a** to a moving positive contrast bar before (black) and after MFA (green) application. **c,** Population data for F-mini-ON RGC spike responses to flashed spots as in **a** (*n* = 3 cells). Points connected by dotted lines are individual cells and bars show mean ± s.e.m. **d,** Same as **c** for peaks of subthreshold voltage responses (*n* = 6 cells). **e,f,** Image of a recorded F-mini-ON RGC (magenta) before and after ablation of 3 of the coupled somas (cyan). **g,** Responses of an F-mini-ON RGC to an ON 120 µm spot at 2000 R*/rod/s before (black) and after ablation (red) of coupled somas. Dashed line is −60 mV. **h,** Responses of a different cell to a moving ON bar at 700 R*/rod/s before (black) and after ablation (red). **i,** Population data for F-mini-ON RGC spike responses to flashed spots as in **c** (*n* = 5 cells). Points connected by dotted lines are individual cells and bars show mean ± s.e.m. **j,** Same as **i** for subthreshold membrane voltage (*n* = 5 cells).

Our second strategy to isolate F-mini-ON RGCs from their coupled network used physical ablation^26^, which had the advantage of increased specificity. Using Alexa Fluor 488 fluorescence under 2P illumination, we were able to visualize the F-mini-OFF RGCs coupled to a targeted F-mini-ON RGC. In the ablation procedure, we recorded light responses in the F-mini-ON RGC before and after destroying the coupled cells by membrane rupture with sharp microelectrodes (Fig. 3h). Ablation eliminated OFF spikes and reduced OFF depolarization to 18 ± 13% of control, while only reducing ON spiking to 77 ± 25% and ON depolarization to 87 ± 18% of control (Fig. 3i-l, *n* = 5 spiking, 6 depolarization). Baseline membrane potential changes following ablation were not significant (−60 ± 3 mV to −58 ± 6 mV, *n* = 6, p = 0.36). Results from both approaches demonstrated that F-mini-ON RGCs receive ON input through canonical synaptic pathways and receive OFF input through a non-canonical pathway involving gap junctions with F-mini-OFF RGCs.

With this knowledge of the different circuits responsible for the ON and OFF parts of the RFs of F-mini-ON RGCs, we returned to our observation of the spatial offset between the ON and OFF RFs in search of a mechanism. The distinctive asymmetric morphology and connection pattern of F-mini RGCs offered an important clue. Coupled somas tended to be positioned dorsal to F-mini-ON RGCs in the ventral and central retina where we performed our measurements (Fig. 4a), consistent with their dorsally directed dendrites in these regions^16^. The dendrites of F-mini-ON and F-mini-OFF RGCs are oriented in opposite directions relative to their somas^16^, and they overlap with each other in stratification only in the central part of the inner plexiform layer (Fig. 1c). Assuming that this shared stratification is the location of the gap junctions, then the main stratification fields of coupled F-mini-OFF RGCs are displaced ventrally from those of the F-mini-ON RGCs to which they are coupled. We created an anatomical RF model using only our measurements of connected soma spatial offsets and soma-to-dendrites morphology (Fig. 4b, see Methods). Model OFF RFs were displaced on average 25 µm ventral from ON RFs (Fig. 4c), similar to the shift we measured in F-mini-ON RGCs (Fig. 1n, −30 ± 13 µm). This result demonstrates that the spatial structure of the F-mini RGC network is sufficient to account for the ON-OFF RF offset we observed in F-mini-ON RGCs.

**Figure 4.**
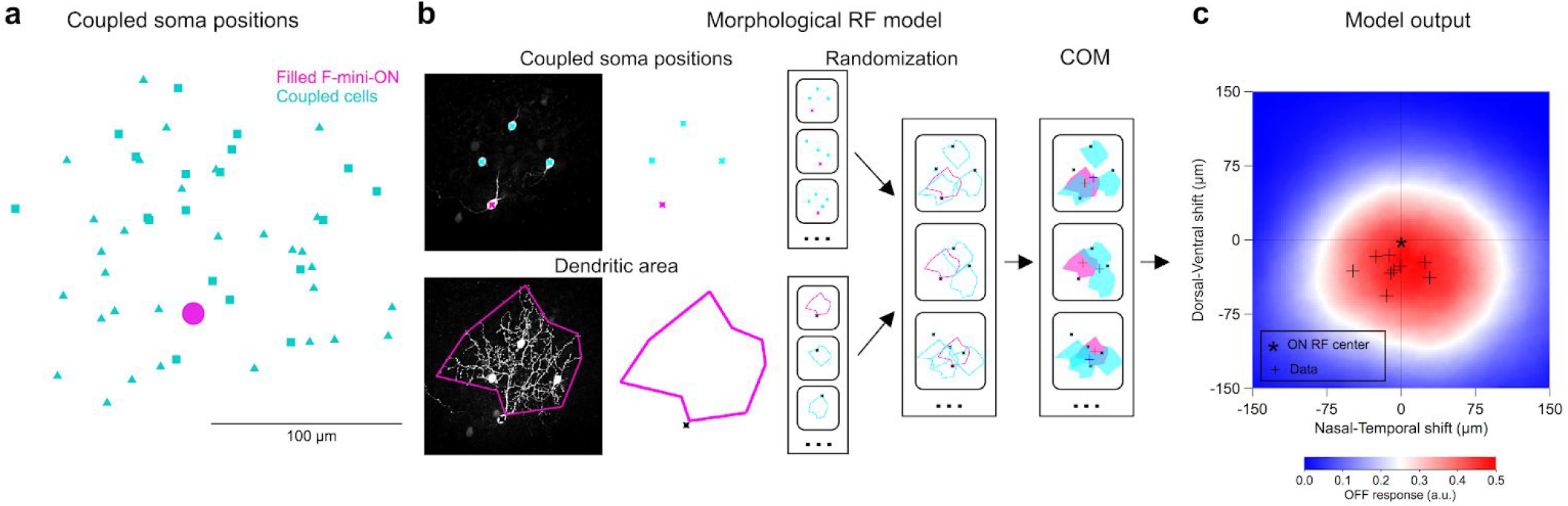
F-mini-ON RGCs RF offset is captured by morphological model. **a,** Locations of labeled somas relative to the injected F-mini-ON RGCs included both confirmed F-mini-OFF RGCs (squares) and RGC somas that were not further characterized (triangles) (*n* = 11 injected cells) Each point represents the position of a gap-junction labeled soma relative to the position of the filled F-mini-ON RGC. Results above suggest that all coupled cells were in fact F-mini-OFF RGCs, but only some of them (squares) were confirmed via electrophysiology or IHC. **b,** RF model diagram (see Methods). Measurements of coupled soma positions from **a** were combined with convex polygon fits to the dendrites of F-mini-ON and F-mini-OFF RGCs, to create a purely anatomical prediction of the center-of-mass (COM) of ON and OFF RFs. The model randomized the selection of the number of coupled somas and coupled soma positions according to our measured distributions (Fig. 2d, and panel **a,** respectively). Polygon fits of the F-mini-ON and F-mini-OFF RGC dendrites were also randomized over our collection of traced images for each cell type. **c,** Results from the dendritic RF model. Data points (black crosses) and format are from Fig 1n.

What visual features are represented by offset ON and OFF sub-regions of the RF? We first tried to answer this question at the level of single RGCs using a model that captured both the spatial structure of the ON-OFF RF center of F-mini-ON RGCs and suppression by their RF surround (Extended Data Fig. 8a,b, see Methods). The model replicated the mild direction selectivity we measured in F-mini-ON RGCs only for slowly moving stimuli (Extended Data Fig. 8c,d), also reported in the initial study of these RGCs^16^. We also found mild orientation selectivity in F-mini-ON RGCs with the presentation of drifting gratings (Extended Data Fig. 8e). In a version of our single-cell model that matched the axis of elongation of the F-mini-ON RFs, we were able to predict a similar degree of orientation selectivity (Extended Data Fig. 8f). However, neither of these properties provides a satisfying explanation for this unique RF structure since other specialized RGCs encode movement direction and orientation with much greater specificity (Extended Data Fig. 8c,e).

Next we constructed a multi-cell model to explore whether offset ON-OFF RFs could provide an encoding benefit in the population that is less apparent at the single-cell level. Specifically, we tested whether a population of RGCs with consistently offset ON-OFF RFs is more precise in representing the position of a dark-light edge than a population of RGCs with either overlapping ON-OFF RFs or with randomly offset RFs (Fig. 5). Our model used (1) overlapped ON and OFF RFs, (2) larger RFs resulting from offsetting the same ON and OFF sub-fields, or (3) RFs with the same overall size, but with offset ON and OFF sub-fields, as we measured in F-mini-ON RGCs (Fig. 5a). The two offset models used either a consistent ventral offset between ON and OFF sub-fields or a random distribution of offset directions, consistent with our measurements from F-mini-ON and control ON-OFF RGCs, respectively (Fig. 1l). All other aspects of the five models were identical. Positions of model cells were generated as a noisy hexagonal grid based on the measured density of F-mini-ON RGCs^13^. Our single-cell model described above generated responses for each RGC using its ON and OFF RF sub-fields and a suppressive surround. Gaussian noise with a magnitude consistent with our spike data from F-mini-ON RGCs, was added to each response. We presented our three models with edge stimuli at various orientations and spatial locations, and we computed the center-of-mass of the model RGC responses to decode edge position (Fig. 5a-c).

**Figure 5.**
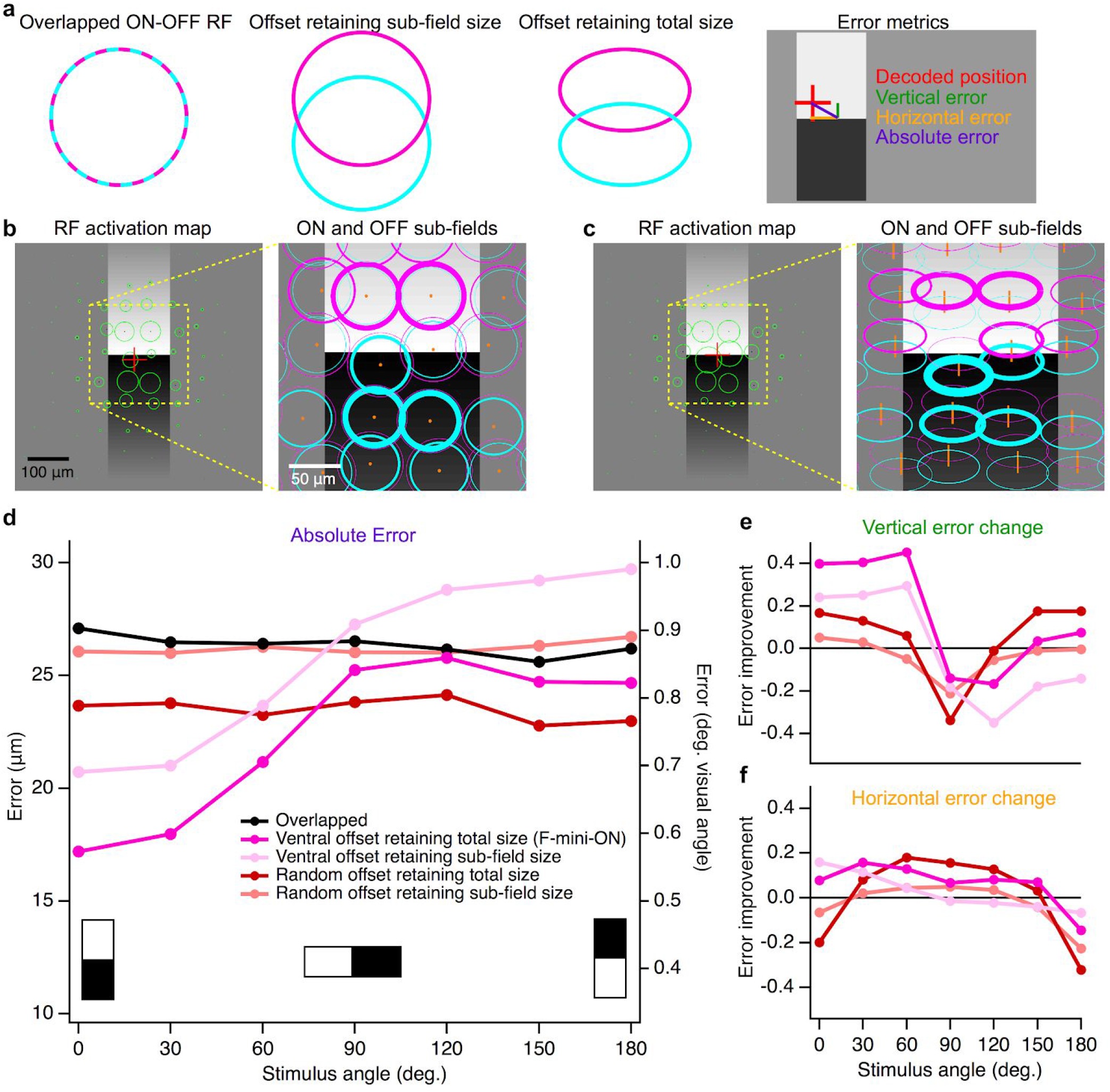
Multi-cell model of object localization shows an advantage of offset ON-OFF RFs. **a,** Schematics of the different RF structures and error metrics. **b,** RF activation map (left) for a single instantiation of the overlapped RF model. The stimulus is the black-white edge. Green circles show the positions of each RGC with the radius of the circle proportional to its response. Red cross indicates the center of mass of the RGC responses. Magnified portion (right) shows the ON (magenta) and OFF (cyan) sub-fields of each RF. Line thickness is proportional to activation of each sub-field. **c,** RF activation map for the same positions and stimulus as in **b,** but for the offset RF model retaining total size. Orange lines in the magnified view (right) connect ON and OFF sub-fields from the same RGC. **d,** Absolute error in decoded edge position for each model. Points are means of 500 iterations of each model with different RGC and stimulus positions. Error bars for s.e.m. are obscured by symbols; s.d. was similar for each point with a mean of 0.42 µm. **e,** Vertical component of the error in **d** shown as a change ratio relative to the overlapped RF result. **f,** Same as **e** for the horizontal component.

Models with consistently offset ON-OFF RFs performed better than those with overlapped or randomly offset RFs in representing the position of an edge stimulus (Fig. 5d-f). Along the (vertical) axis of separation of the RF sub-fields, F-mini-ON RGC model was 40% better than the overlapped RF model at representing the position of horizontal edges (Fig. 5d), particularly the vertical component perpendicular to the edge (Fig. 5e). This improvement came with no decrease in horizontal position decoding performance (Fig. 5f), but with a tradeoff in performance for edges having a contrast offset perpendicular or opposed to the RF offset. The F-mini-ON RGC population model was able to represent the position of an edge with precision down to 0.6 degrees of visual angle (Fig. 5f), less than 12% of the (2 sigma) RF diameter of a single F-mini-ON RGC. The improvement of the offset models relative to the overlapped RF model was robust across a broad range of cell densities, RF sizes, and noise amplitudes (Extended Data Fig. 9).

## Discussion

Collectively, our results demonstrate that the F-mini-ON RGC mixes a canonical ON input via chemical synapses with a non-canonical OFF input via gap junctions with F-mini-OFF RGCs to create an RF with spatially offset ON and OFF regions. The offset is consistent with the asymmetric morphology and connectivity of F-mini RGCs (Fig. 4). Our multi-cell model shows that the ON-OFF RF offset can improve the precision with which F-mini-ON RGC populations represent the position of an edge (Fig. 5). A causal link between this proposed role in encoding edge location and a specific behavior will require selective genetic access to F-mini RGCs. With advances in molecular profiling of RGCs^14^, the tools for such a study are on the horizon.

Recent work has shown that mice use vision to capture small, quickly moving prey objects, like crickets^27,28^, so perhaps precise edge localization with F-mini-ON RGCs plays a role in this behavior. Since the RF offset in F-mini-ON RGCs is along the vertical axis, our model showed enhanced object localization preferably along this axis (Fig. 5). Future behavioral analyses could determine whether mice show more precise object localization along the vertical than the horizontal axis of the visual field. Since rodents use compensatory eye movements to maintain the orientation of their eyes relative to the horizon^29^, a potential advantage of precise localization along the vertical axis is that it would provide information about changes in distance: approaching objects have increasing space between their dorsal and ventral edges with time, while receding objects have decreasing space. The direction of the ON-OFF asymmetry is also interesting as it relates to rodent visual ethology. Dark-below-light edges in retinal space were represented best by our model F-mini-ON RGC population (Fig. 5d). This corresponds to dark-above-light edges in visual space, consistent with the special behavioral relevance of “looming” dark objects in the upper visual field^30^.

Heterotypic RGC coupling was also recently identified in the guinea pig using multi-electrode array recordings and morphological tracing with Neurobiotin^31^. The coupled RGCs in the guinea pig study were both ON sustained cells – one of them was the ON alpha – so heterotypic coupling in that system does not mix ON and OFF signals. Nonetheless, the discovery of heterotypic RGC coupling in two different circuits in two different species suggests that it might be a conserved motif in the mammalian retina, augmenting our understanding of the organization of information in RGC populations.

Direct electrical coupling between RGCs has long been known to be important for synchronizing spikes on the timescale of several milliseconds^32–34^, and tight spike synchrony has been shown to enhance transmission at retino-geniculate synapses^35^. Could synchronous firing of coupled F-mini-ON and F-mini-OFF RGCs encode different features of the visual scene than firing of either cell alone? Synchronous firing among RGCs has been proposed as a mechanism for motion anticipation^7,19^, improving the fidelity of the population code for direction selectivity^36^, and binding objects across space^18^. Whether spike synchrony the F-mini RGC network participates in these or other computations is an attractive question for future work.

The dynamics of the F-mini RGC network are another target for future studies. The strength of gap junctions in the retina, including those between RGCs, can be altered by neuromodulators^17,20,37^. If the F-mini RGC network is modulated, this could change its function with sensory or behavioral context. Finally, comparisons with other species will provide information about the evolution and function of this particular heterotypically-connected RGC circuit. FoxP2 also labels a subset of RGCs in the ferret^38^ and macaque retinas^16^, so it is possible that the F-mini RGC circuit has a homolog in humans.

## Methods

### Animals and electrophysiology

The retinas of wild-type mice (C57BL/6J, Jackson Labs) were used for all experiments. The mice were dark adapted overnight and euthanized by cervical dislocation in accordance with all animal care standards provided by Northwestern University’s Institutional Animal Care and Use Committee. Lighting in animal facilities was kept on a 14/10 hour cycle, with lights on at 6:00 am. Typical retina in vitro times were 12:00 pm through 7:00 pm. Mice of either sex, and ages P30-P90 were used; no differences in results were observed with sex or age. Eyes were dissected in oxygenated Ames medium at 32°C. Dissections were performed in complete darkness using infrared (IR, 900 nm) illumination and photo converters. In the experimental rig, retinas were mounted in a shallow dish, below a microscope objective and above a digital projector, in oxygenated Ames medium at 32°C at a flow rate of 10 ml/min. Two glass electrodes on headstage amplifiers were mounted on micromanipulators on either side. Cell-attached and current clamp experiments were performed as previously described^26^.

### Microscopy

Two microscopes were used to visualize cell morphology. The *in vitro* microscope, a Scientifica SliceScope Pro 6000, used 980 nm illumination from a SpectraPhysics MaiTai Laser steered by a galvo scanner for 2P excitation and IR visualization. Software was SciScan version 1.5 by Scientifica in LabVIEW by National Instruments. Dyes for two-photon visualization were Alexa Fluor 488 and 568, the latter of which was found to be not gap junction permeable and was used for single-cell image isolation. Microscopy was continued on fixed retinas, which were stained with antibodies and fluorescent dyes for immunohistochemistry. The fixed-tissue microscope was a Nikon A1R confocal microscope with a 1.0 NA 40x oil immersion objective at the Northwestern Center for Advanced Microscopy.

### Light stimulation

Spatio-temporal light patterns were focused on the photoreceptors of the *in vitro* retina.The light patterns were generated by a computerized digital display, a DLP Lightcrafter 4500 from Texas Instruments, illuminated by a blue LED at 457nm (peak wavelength after optics), integrated by EKB Technologies Ltd. It had a resolution of 1140 × 912 pixels, operating at 60 Hz, frames modulated to 8 bits intensity depth. Overall light intensity was modulated using neutral density filters (Thorlabs) and calibrated regularly. Measured intensity values were converted to rhodopsin isomerizations per rod per second (R*/rod/s). Light stimuli were generated by MATLAB software packages: Schwartz Lab protocols (https://github.com/Schwartz-AlaLaurila-Labs/sa-labs-extension) interfacing with the Symphony 2 Data Acquisition System (https://symphony-das.github.io) drawing via Stage (http://stage-vss.github.io) and OpenGL to the screen. The microscope’s condenser brought the projector image to focus at the photoreceptor outer segments at a scale of 1.38 µm/pixel. Stimuli included circles (30 - 1200 µm in diameter) flashed with positive or negative contrast for 1.0 s, moving bars (180 - 1200 µm width by 600 - 1200 µm length, at 250 - 2000 μm/s speed), and small spots for RF mapping (see below). Stimulus for the light stimulation cross correlation analysis was a randomly moving light spot 80 µm diameter, 30 sec duration. The random motion path was a zero-centered 2D gaussian noise signal, lowpass filtered at 3 Hz cutoff frequency. Light stimuli were presented from darkness (< 2 rod isomerizations R*/rod/s) to a level of 200 R*/rod/s unless otherwise noted, in order to preserve the sensitivity of the retina.

### Receptive field mapping

Current-clamp recordings allowed us to measure subthreshold voltage responses with small spots to obtain high resolution RF maps. Visual stimulus spots were circles of positive and negative 100% contrast on a background of 1000 R*/rod/sec, with a diameter of 40 µm. A triangular grid of 30 µm spacing was used for F-mini RGCs. Larger spacing and spot sizes up to 80 µm were used for RGCs with larger RFs and lower sensitivity to small spots. Voltage responses to individual spots were separated and peak values were averaged to generate a value for each position. These values were displayed on the grid locations to create a 2D RF strength map. The center of mass of the map area above the 80-85th percentile of response strength was used to generate offset vectors for comparison to the model. The RF overlap index (Fig. 1k) uses the proportion of overlap relative to the total of the RF area within the 80th percentile of response strength. Control ON-OFF RGCs were of types UHD, HD2, and On-Off DS, which exhibit similar ON-OFF transient responses to small spots.

### Cell identification

Somas in the ganglion cell layer were surveyed in cell-attached mode using a set of basic light stimuli: flashing contrast steps in 160 µm spots, spots of varying sizes, and moving bars. F-mini RGCs could be identified by their characteristic responses to these stimuli. Once an F-mini RGC was identified, it was dye filled and recorded to verify type and collect data. Pairs were generally done sequentially: (1) Identify F-mini-ON/OFF RGC by its light response, (2) fill with Alexa Fluor 488, (3) use dye to find coupled F-mini-OFF/ON RGCs. Sequential filling of coupled F-mini RGCs with Alexa Fluor 488 allowed for identification of large networks of > 10 cells. Neurobiotin in a single F-mini RGC labels second order connected cells more dimly than first order ones, in fixed tissue, allowing for similarly large network identification.

### Pharmacology

Meclofenamic acid (MFA) was bath-applied at 100 μM to block gap junctions (Figs. 2, 3, Extended Data Figs. 4, 5)^23^. Electrophysiology results in MFA conditions were 5 through 45 minutes during application. MFA washes out incompletely, so no post-application data is reported.

### Ablation

Neighbor ablation is a physical technique for neuronal inactivation where a micropipette is used to rupture the cellular and nuclear membranes of the dye-illuminated somas^26^. This causes the membrane of the entire cell to dissociate, and stops it from having a membrane potential or transmitting and receiving signals within the dendrites. Neuron death was confirmed by lack of Alexa Fluor 488 in 2P imaging. Some responses continued for one stimulus epoch before being silenced. Where incomplete network ablation occurred, changes to response properties were partial. These data were not used further.

### Immunohistochemistry

Target neurons were filled with Neurobiotin from Vector Laboratories SP-1150 3% w/v and 280 mOsm in our standard potassium aspartate internal solution^26^. For whole mount immunohistochemistry (IHC), post-*in vitro* retinas were fixed in 4% paraformaldehyde for 15 minutes, then rinsed in phosphate buffer. Primary antibodies for cell typology were FoxP1 to guinea pig from Prof. Bennett Novich^39^ and FoxP2 to rabbit from Millipore (ABE73). Primary antibodies for connexin typology were Cx30.2 from Invitrogen (40-7400), Cx36 to rabbit from Invitrogen (51-6200), and Cx45 to mouse from Invitrogen (41-5800). Retinas were soaked with primary antibodies in normal donkey serum (NDS) + Triton-X for 3 days at 4°C. Secondary antibodies/fluorophores were (for typology) donkey anti-rabbit Alexa Fluor 568 from Life Technologies (A10042), goat anti-guinea pig Alexa Fluor 647 from abcam (ab150187), (for connexins) donkey anti-rabbit Alexa Fluor 568 from abcam (ab175470), donkey anti-rabbit Alexa Fluor 647 from Jackson ImmunoResearch (711-605-152), donkey anti-mouse Cyanine Cy3 from Jackson ImmunoResearch (715-165-150), and Streptavidin DyLight Conjugate 488 from Thermo Science (21832). Retinas were soaked in secondary antibodies in NDS + Triton-X for one day at 4°C. Retinas were glass slide mounted in Fluoromount Aqueous mounting medium from Sigma-Aldrich (F4680) and stored at 4°C.

### Morphological receptive field model

Receptive fields relative to soma location of F-mini-ON and F-mini-OFF RGCs were estimated using the area between the tips dendritic fields (*n* = 38 F-mini-ON, *n* = 12 F-mini-OFF). These were traced manually using 2P or confocal image stacks using the SimpleNeurite Tracer plugin in ImageJ. Locations of F-mini RGCs in coupled networks were traced from images of dye-filled somas to create maps of soma locations (*n* = 11). Random combinations of network soma locations, F-mini-ON dendrite offsets, and F-mini-OFF dendrite offsets were generated 1000 times and averaged to generate a mean OFF RF relative to ON RF (Fig. 4d). The model ignores any possible interdependence of F-mini-ON and F-mini-OFF RGC dendritic fields (meta-mosaics), and assumes that F-mini-OFF RGCs receive OFF input via bipolar cells at their dendritic tips.

### Single-cell RF model

A computational model (Extended Data Fig. 8a) was used to generate single cell responses to an edge, moving bar, and drifting grating stimulus. The model simulated four pathways of input to a single RGC: ON and OFF, excitation and inhibition for each. Excitation was modeled as a small 2D gaussian of direct excitation with a larger 2D gaussian subtracted to model presynaptic inhibition. Direct inhibition was modeled as a larger 2D gaussian. The visual stimulus was multiplied by the spatial RFs, then those signals were integrated across space and temporally filtered by convolution. Temporal filter kernels were parameterized curves with values extracted from typical F-mini-ON voltage-clamp light-step responses, over a 3.0 sec simulation time. A semi-rectifying nonlinearity was applied to each ON and OFF sub-field, then the responses were summed. OFF delay relative to ON was estimated from spike latencies. RF sizes and surround strength was adjusted manually to match F-mini-ON responses to spots of multiple sizes. This model is meant to explore RF map concepts analytically over many variables, and is not meant to precisely emulate recorded RGCs.

### Multi-cell decoding model

The multi-cell model composed responses from many single-cell RF instances and decoded them using their center-of-mass. Gaussian noise was added to each modeled cell’s response. The decoded location of the stimulus was compared to the true location to generate an error value. Trials of randomly placed cells and stimulus were used, with 500 for each parameter configuration. Cells RF centers were laid out on an equilateral triangular grid with a gaussian jitter of 10 µm sigma. Receptive field strengths were integral normalized across shape parameters. Comparisons across parameters used a density of 250 RGCs/mm^2^, noise of 2 (a.u., but similar to spiking output), and a stimulus angle of 0 (horizontal, ON upper). RGC responses fell to baseline outside of the model region, which had an area of 0.36 mm^2^ with 600 µm side length. The stimulus was placed randomly uniformly within the central 300 by 300 µm square region. The edge stimulus was a 150 µm length edge of positive and negative 100% contrast, falling off in a linear gradient above and below the edge for 150 µm. Rousso et al^16^ found a range of densities of between 100 and 350 RGCs/mm^2^ for F-mini-ON RGCs. Eyewire Museum’s patch of retina, the E2198 dataset^13^, has a density for F-mini-ON RGCs (type 63) of approximately 240 RGCs/mm^2^.

### Analysis and statistics

Analysis was performed with a custom MATLAB software package. Figures were generated in Igor 8.0 from Wavemetrics. Stimulus code is available at https://github.com/Schwartz-AlaLaurila-Labs/sa-labs-extension. Subthreshold membrane potential measurements (Fig. 3d,j) used a spike-removal lowpass filter of 100 Hz cutoff frequency. All data are reported as mean ± s.d. unless otherwise noted. Comparisons for statistical significance were performed with a paired or unpaired Student’s t-test, as appropriate, unless otherwise noted. Direction and orientation selectivity indices were calculated as the normalized magnitude of the vector sum of the responses across directions or orientations. For measurements of connexin overlap, RGCs were traced using Simple Neurite Tracer in Fiji software. Receptive field ellipticity was measured using the 80th percentile response contour, finding the longest line contained within that (*a*), then the longest such line perpendicular to that line (*b*). The ellipticity is then lengths (*a* − *b*)/*a*.

## Author contributions

S.C. and G.W.S performed experiments. S.C. analyzed data and constructed models. S.C. and G.W.S. designed research and wrote the paper.

## Acknowledgements

We thank Prof. Bennett Novich^39^ for generously providing the FoxP1 antibody.

Imaging work was performed at the Northwestern University Center for Advanced Microscopy, generously supported by NCI CCSG P30 CA060553 awarded to the Robert H Lurie Comprehensive Cancer Center. Multiphoton microscopy was performed on a Nikon A1R multiphoton microscope, acquired through the support of NIH 1S10OD010398-01.

This work was supported by NIH National Eye Institute F31 EY029593, NIH National Eye Institute T32 EY025202, and NIH DP2 EY026770.

