## Extended data for "An offset ON-OFF receptive field is created by gap junctions between distinct types of retinal ganglion cells"

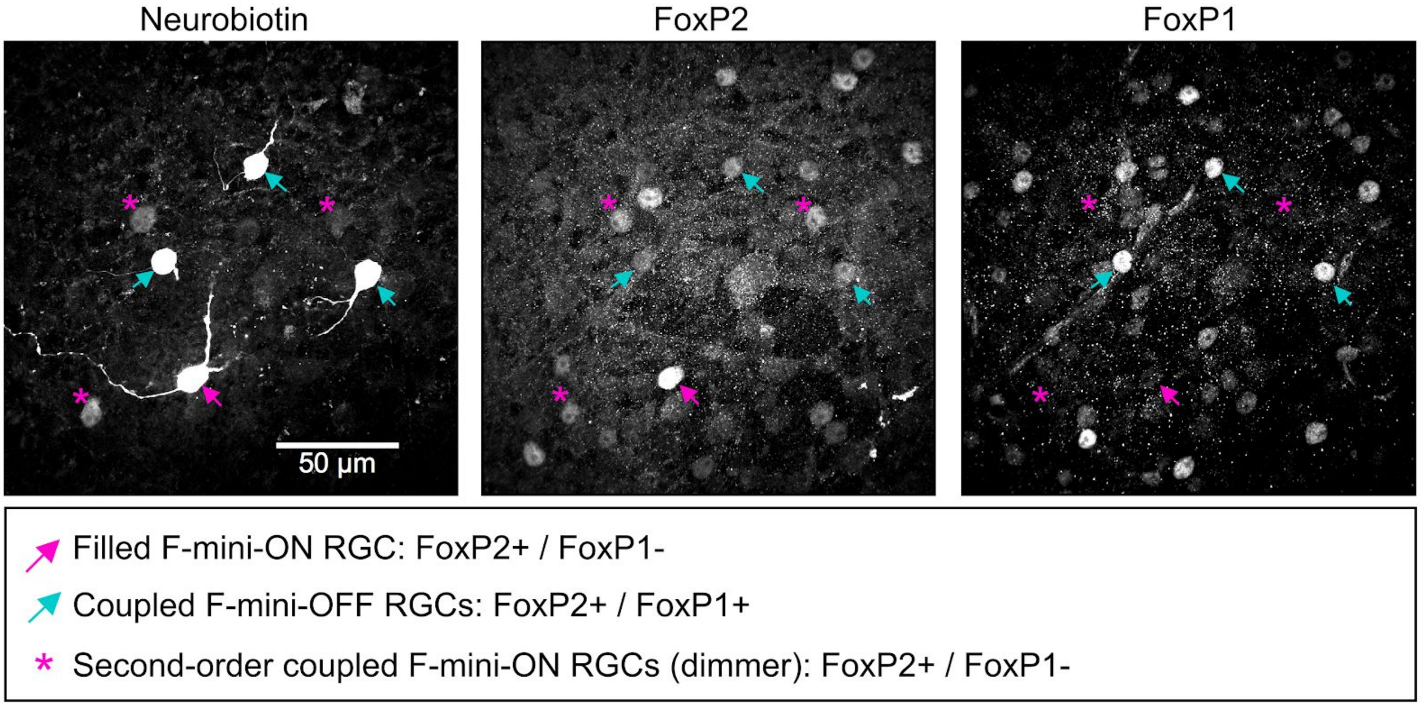


Extended Data Figure 1 **Coupled cells are immunoreactive for F-mini RGC markers.**

Images of the ganglion cell layer in a patch of retina in which a single F-mini-ON RGC was filled with Neurobiotin (magenta arrowhead). Left panel shows the Neurobiotin channel, with three brightly labelled coupled cells (white arrowheads) and three dimly labelled cells that likely represent second-order connections (magenta asterisks). Middle panel shows the same region with immunoreactivity for FoxP1, which labels F-mini-OFF RGCs, but does not label F-mini-ON RGCs[1]. Right panel shows immunoreactivity for FoxP2, which labels both F-mini RGC types. This experiment was performed on five F-mini RGC networks in four retinas: four F-mini-ON RGCs and one F-mini-OFF RGC injected. Three networks were stained for FoxP2 and FoxP1; two networks for FoxP2 only. Neurobiotin labeled 9.0 ± 6.4 somas per retina, and was found in varying amounts in neurons; indicating first and second order connectivity. FoxP2 was present in 43 of 45 RGCs that were labeled with Neurobiotin. Coupled cells from these networks that could be morphologically identified by using the visible primary dendrites, and all showed the expected patterns of FoxP1 expression. 8/8 F-mini-ON RGCs were FoxP1 negative and 14/14 F-mini-OFF RGCs were FoxP1 positive.

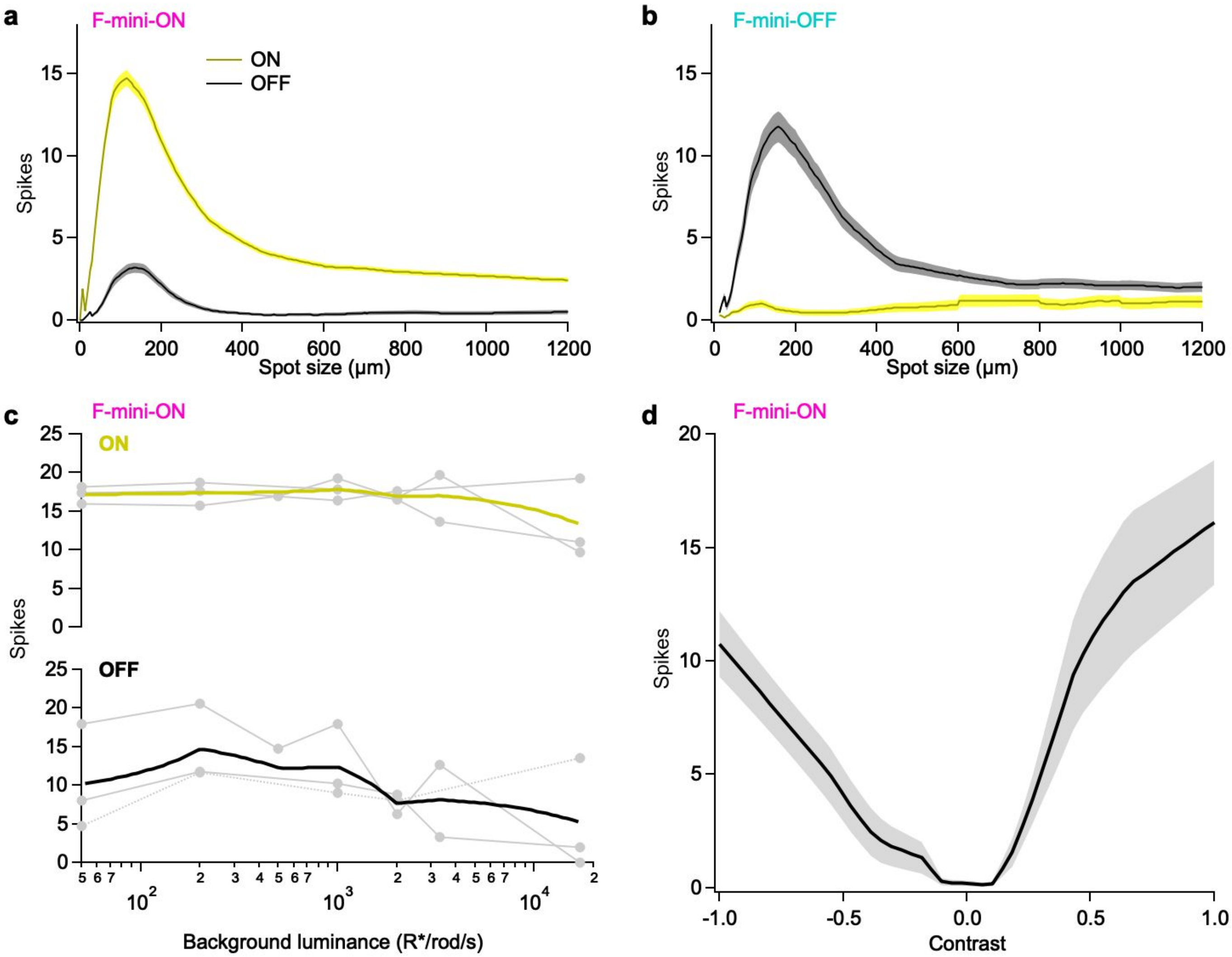


Extended Data Figure 2 **F-mini RGCs have a range of ON-OFF responses.**

**a,** Population data of spike counts in F-mini-ON RGCs responding to positive contrast spots of varying diameters. Onset responses in yellow; offset responses in black. Shaded region is SEM across cells (*n* = 172). Onset responses were well fit by a difference-of-Gaussians model (not shown). Parameters for the model were as follows: SD of center Gaussian = 76 μm, SD of surround Gaussian = 220 μm, center:surround weight ratio = 1.06 **b,** Same as **a** for F-mini-OFF RGCs (*n* = 85). **c,** Spiking responses of F-mini-ON RGCs to flashed spots at varying mean luminance, showing the variations in ON and OFF responses with light level (*n* = 3, spot diameter = 130 μm). **d,** Spiking responses of F-mini-ON RGCs to onset of flashed spots of varying contrast from the background luminance. Shaded region is SD across cells (*n* = 8).

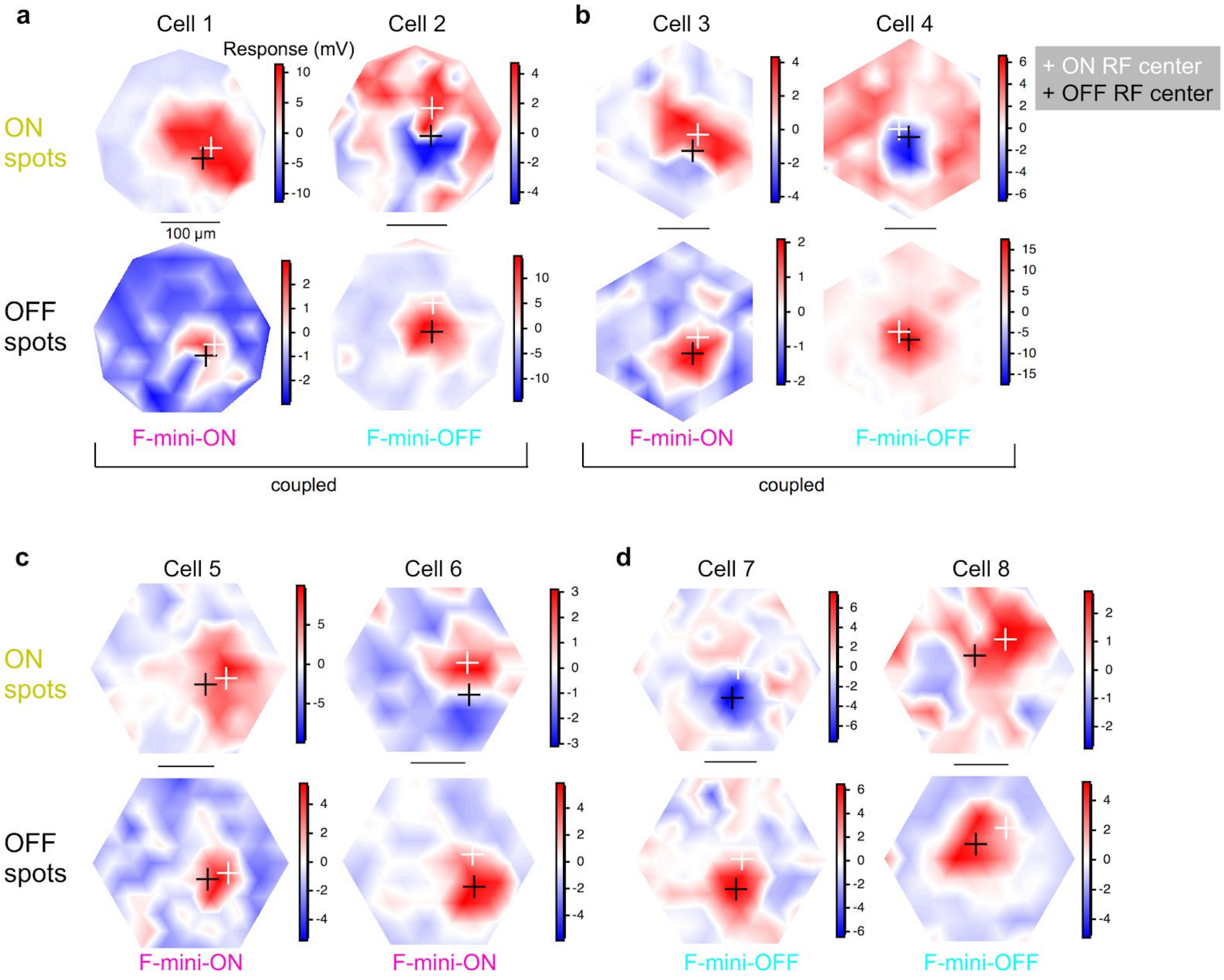


Extended Data Figure 3 **Example RF maps from F-mini-ON and F-mini-OFF RGCs**
Receptive field maps of peak response to 40 µm flashed spots over the RF area, averaged over 2 or 3 repeats. **a,** A GJ coupled F-mini-ON and F-mini-OFF recorded simultaneously. **b,** Another such RGC pair. **c,** Two unconnected F-mini-ON RGCs. **d,** Two unconnected F-mini-OFF RGCs. On all plots, the cross markers are at the center of mass of responses over the 80th percentile (ON, white; OFF, black). Color scale is in mV change from baseline. All scale bars are 100 µm.

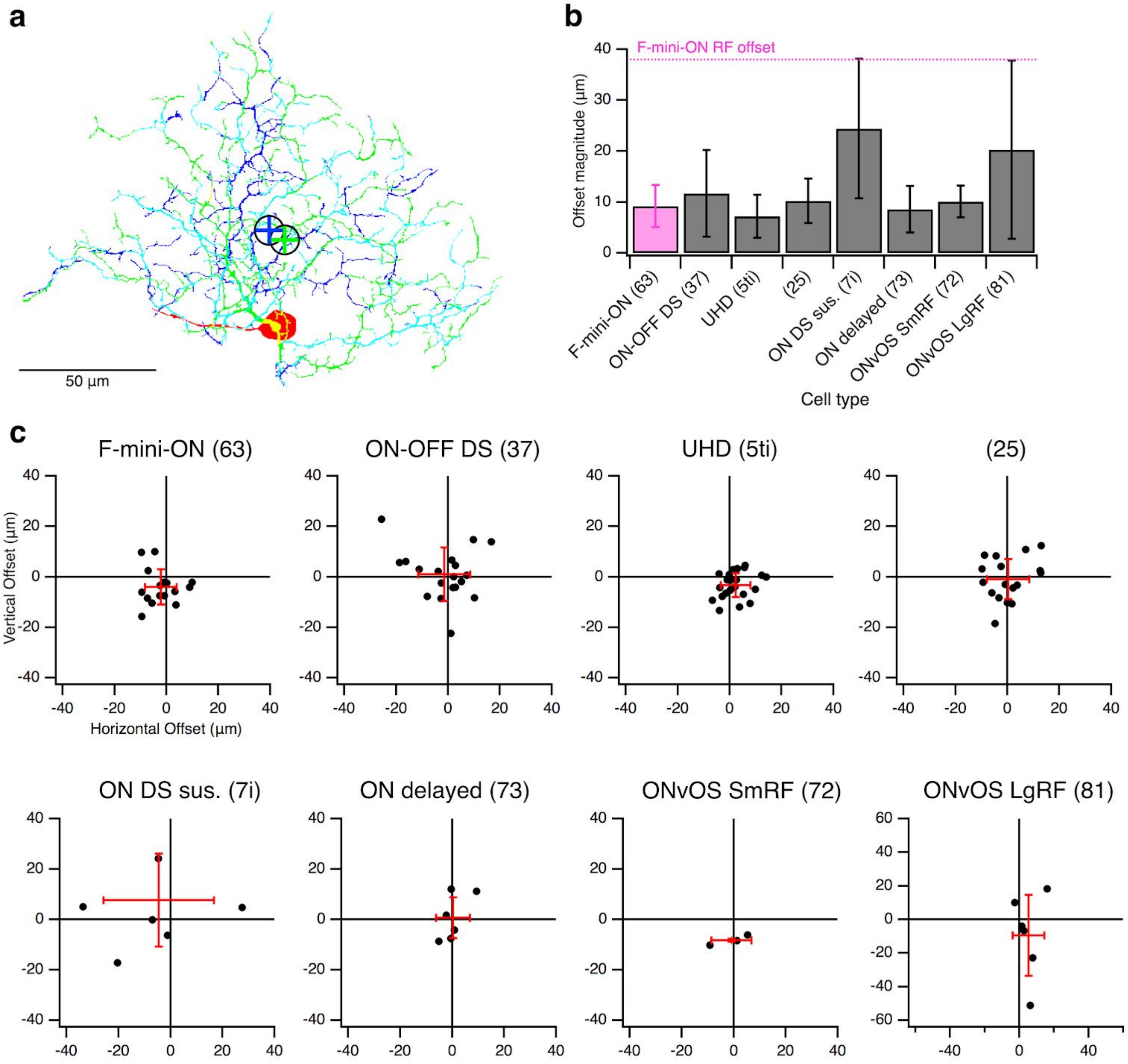


Extended Data Figure 4 **Alignment between ON and OFF strata of bistratified RGCs.**
**a,** Example projection image of an F-mini-ON RGC from Eyewire[2], dendrites colored by stratification: green proximal/inner, blue distal/outer, cyan between, red soma and axon. Centers of mass (COM) of inner and outer strata are marked by circled crosses. **b,** Dendritic offset distance by cell type, mean with SD. Magenta line is at 28 µm, the mean RF offset found in F-mini-ON RGCs. **c,** Offset values in µm from each bistratified RGC in Eyewire by type, listed with corresponding functional type name. Offsets are measured as a vector from proximal/inner COM to distal/outer COM, which in most RGCs is ON to OFF dendrites. Mean and SD are shown by red crosses. All figure data is from the Eyewire dataset[2], exported via the Eyewire Museum mesh tool. Meshes were flattened and offset by eye.

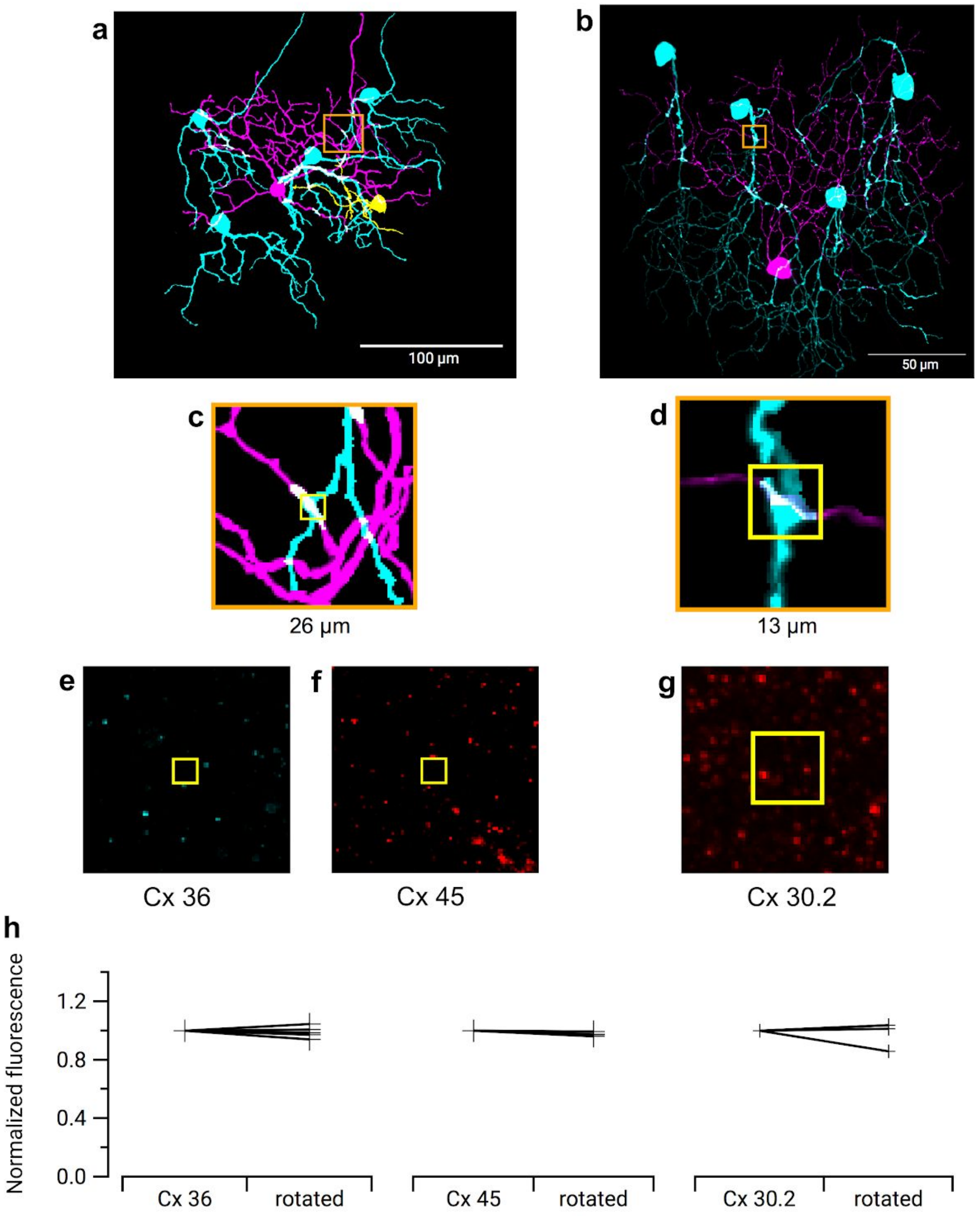


Extended Data Figure 5 **Immunohistochemistry for three types of Connexin at RGC contact points shows negative results**

Three connexins were evaluated for presence at the regions of contact between an F-mini-ON and multiple F-mini-OFF RGCs. **a,b,** Full depth maximum intensity projection images of a Neurobiotin-filled F-mini-ON RGC (magenta) and the connected F-mini-OFF RGCs (cyan). Tracing, segmentation, and masking were performed manually. Image brightness was scaled separately by cell type for illustration here but not for analysis. **c,d** Thin projection images of an example RGC crossing point with yellow square for spatial reference. Stack depth is 3.5 µm. **e-g,** The same region and depth as in **c,d,** showing the IHC channels for the three connexin proteins. **h,** Quantification of overlap between connexin images and RGC contact region masks. Values are similar before and after a 90 degree rotation of the connexin image. Points mark the overlap of the single F-mini-ON RGC with each F-mini-OFF RGC in the image.

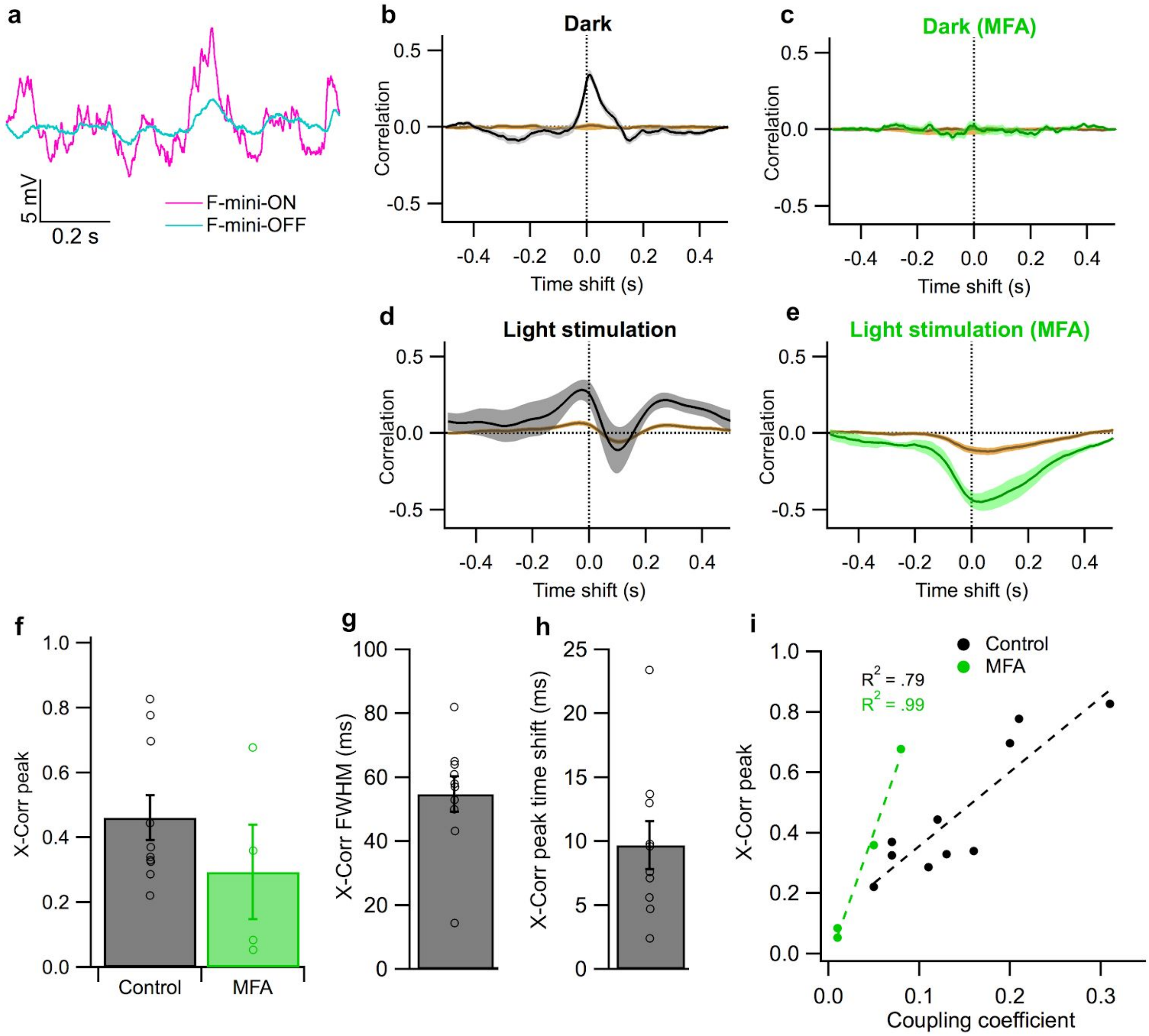


Extended Data Figure 6 **Noise correlations between F-mini-ON and F-mini-OFF RGCs.**

**a,** Traces from a simultaneously recorded pair of F-mini-ON (magenta) and F-mini-OFF (cyan) RGCs in current clamp in darkness (no stimulus). **b-e**. Example cross correlation of the simultaneous voltage from the cells in **a**. Brown trace is for shuffled trials. Shaded regions are SEM across trials. Time shift is F-mini-ON - F-mini-OFF (positive values are F-mini-ON earlier). **b,** Results in darkness. **c,** Results in darkness in the presence of MFA. **d,** Results under randomly moving object light stimulation. **e,** Results under the same light stimulation in the presence of MFA. **f,** Population data showing peak cross-correlation in control and in MFA. Values in MFA are significantly lower than corresponding values in control ($p = 0.007$, one-sided Student's t-test). **g,** Full width at half max and **h,** time shift (right) of cross correlation peak in control conditions. Error bars in **f-h** are SEM across cell pairs and points are each cell pair. **i,** Relationship between cross-correlation peak and coupling coefficient in darkness measured from current injections as in Fig. 2e-h.

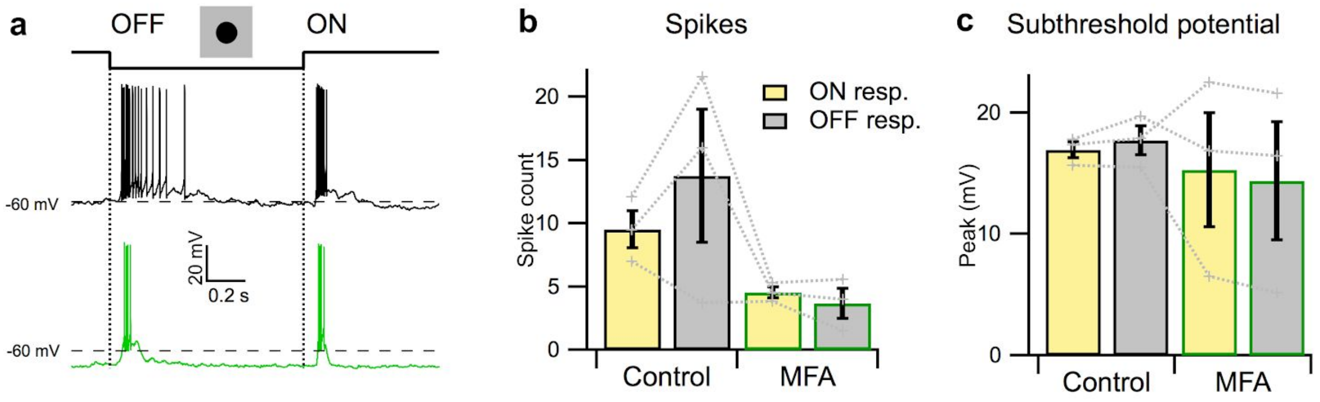


Extended Data Figure 7 **MFA does not selectively eliminate OFF responses in non-F-mini RGCs.**
**a,** Example of an ON-OFF direction selective RGC responding to the onset and offset of a dark spot from a mean luminance of 2000 R*/rod/s in control conditions (black) and in MFA (green). **b,** Population data of spike counts and **c,** subthreshold potential responses to an OFF light step as in **a** for 3 ON-OFF DS RGCS. Baseline voltage level shift mean in control RGCs was -59.9 to -61.8 mV (*n* = 3).

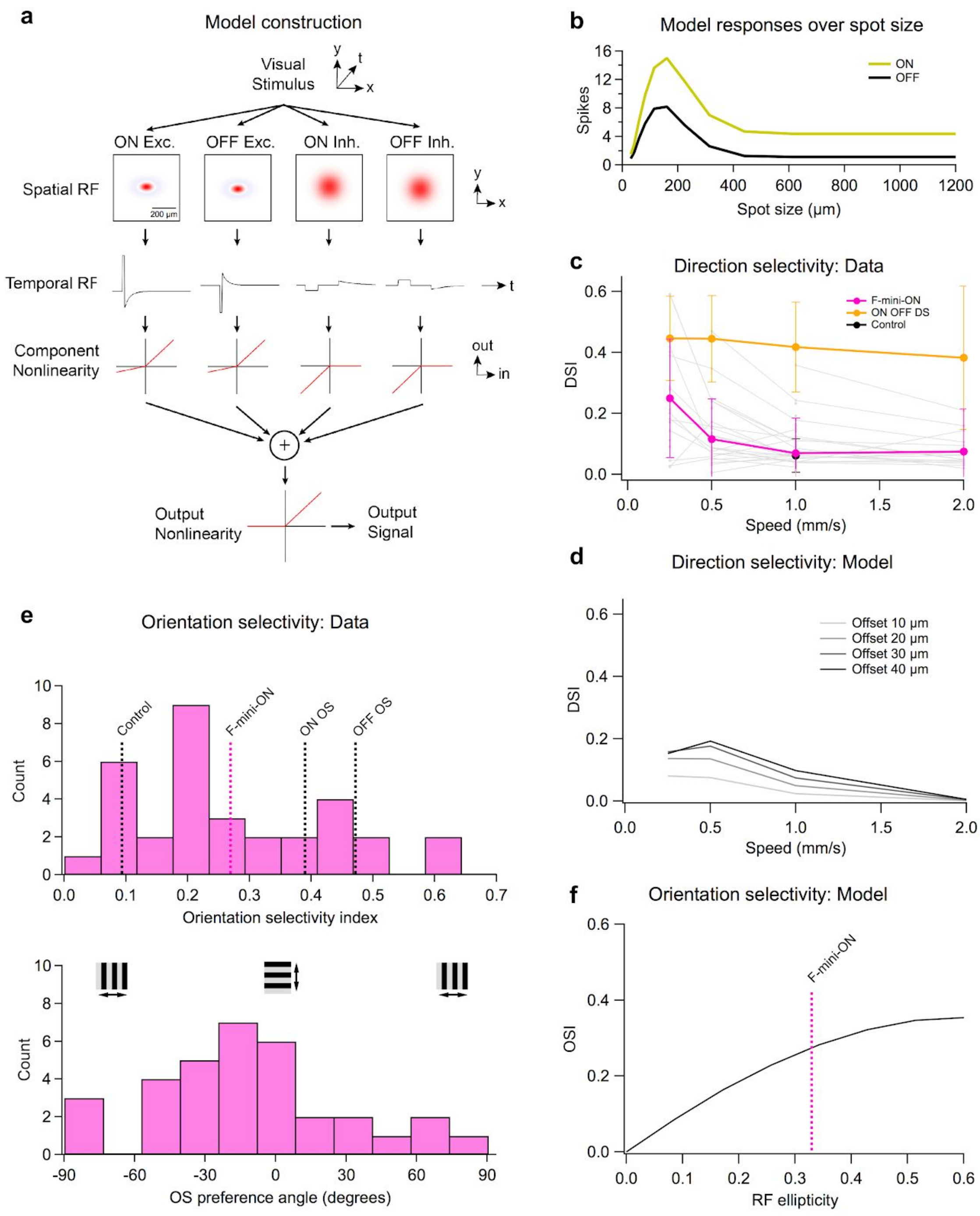


Extended Data Figure 8 **A single cell model generates responses similar to those observed in F-mini-ON RGCs**

**a,** Diagram of single cell receptive field offset model showing the parameters for each of four RGC input component pathways. **b,** Responses of the model to flashed spots of varying sizes showing a qualitative match of surround properties to F-mini-ON RGCs as seen in Extended Data Figure 2a. **c,** Measured direction selectivity in F-mini-ON and ON-OFF DS RGCs, varying over speed. Individual F-mini-ON RGCs are shown in gray (*n* = 103 F-mini-ON and *n* = 279 ON-OFF DS). **d,** Model response DSI over object speed showing similar DSI magnitude and low-speed preference properties to measured responses. **e,** (upper) Orientation selectivity of the population of F-mini-ON RGCs. Dashed lines are published means for OS and control RGCs[3,4]. (lower) Distribution of OS preference angle.

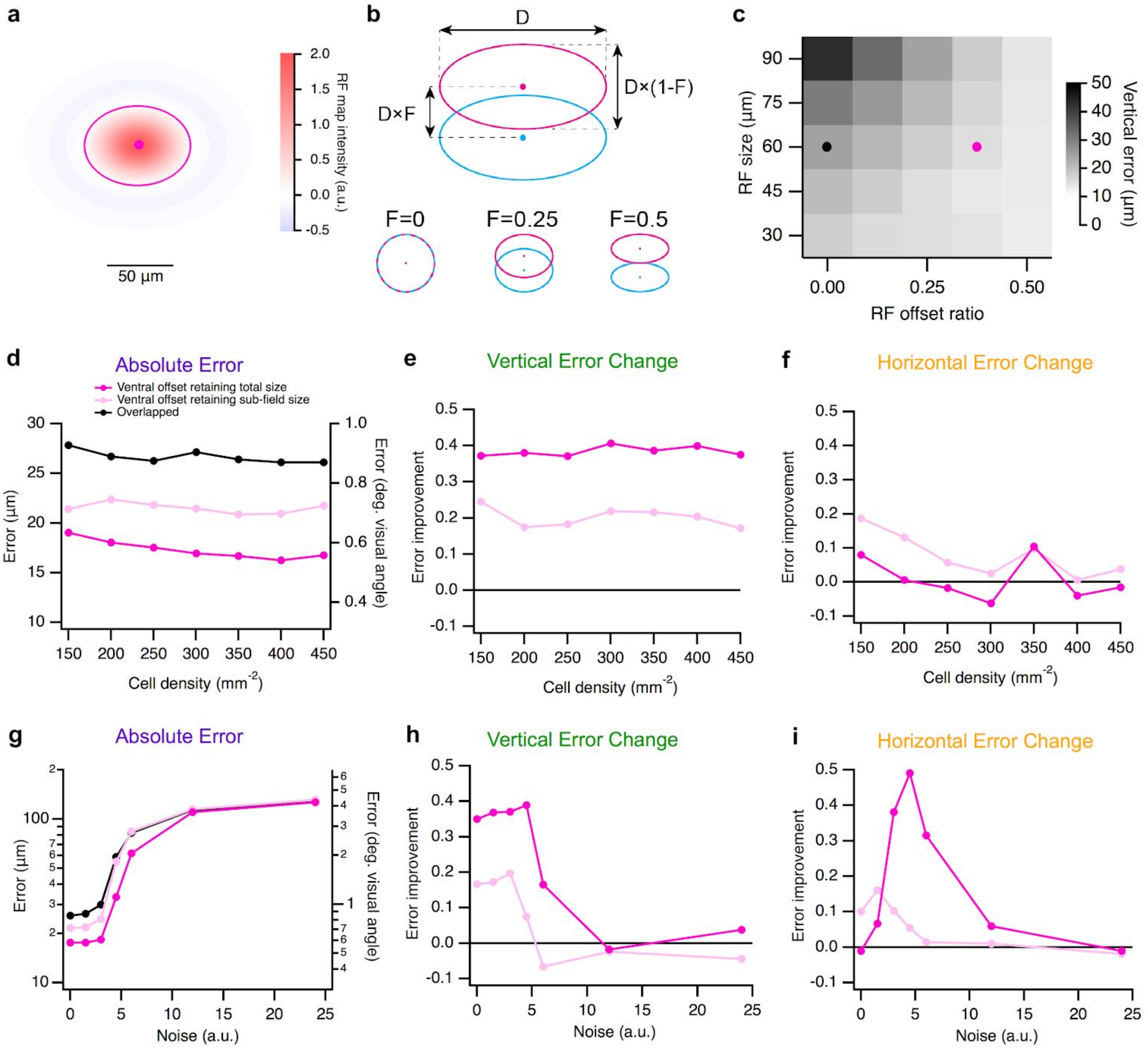


Extended Data Figure 9 **Multi-cell model results are robust over several parameters**

**a,** Illustration of the difference of gaussians RF map used in the single cell model, with an ellipse at the central 2σ contour. **b,** Diagram of RF offset and scaling properties in the model: the diameter (D) and the offset ratio (F) between ON (magenta) and OFF (cyan) sub-fields. **c,** Heatmap of vertical position error (for horizontally oriented stimuli) across models with a range of RF size (D) and RF offset ratio (F). Black and magenta points are the parameters used in the following panels and those in Fig. 5d-f. **d,** Absolute error, **e,** vertical error change ratio, and **f,** horizontal error change ratio for the three RF models across a range of cell density. **g,** Absolute error, **h,** vertical error change, and **i,** horizontal error change ratio for the three RF models across a range of noise values.

**Extended Data References**


1. Rousso, D. L. *et al.* Two Pairs of ON and OFF Retinal Ganglion Cells Are Defined by Intersectional Patterns of Transcription Factor Expression. *Cell Rep.* **15**, 1930–1944 (2016).
2. Bae, J. A. *et al.* Digital Museum of Retinal Ganglion Cells with Dense Anatomy and Physiology. *Cell* **173**, 1293–1306.e19 (2018).
3. Nath, A. & Schwartz, G. W. Cardinal Orientation Selectivity Is Represented by Two Distinct Ganglion Cell Types in Mouse Retina. *J. Neurosci.* **36**, 3208–3221 (2016).
4. Nath, A. & Schwartz, G. W. Electrical synapses convey orientation selectivity in the mouse retina. *Nat. Commun.* **8**, 2025 (2017).
